# SLFN11 captures cancer-immunity interactions associated with platinum sensitivity in ovarian cancer

**DOI:** 10.1101/2020.05.22.110593

**Authors:** Claudia Winkler, Domenico Ferraioli, Anna Garuti, Federica Grillo, Jaime Rodriguez-Canales, Lorenzo Ferrando, Nicolas Chopin, Isabelle Ray-Coquard, Davide Bedognetti, Alberto Ballestrero, Elisabetta Leo, Gabriele Zoppoli

## Abstract

Large independent analyses on cancer cell lines followed by functional studies have identified Schlafen 11 (SLFN11), a putative DNA/RNA helicase, as the strongest predictor of sensitivity to DNA-damaging agents. However, its role as a prognostic biomarker is undefined, partially due to the lack of validated methods to score SLFN11 in human tissues. Here, we implemented a pipeline to quantify SLFN11 in human cancer samples. By analyzing a cohort of high-grade serous ovarian carcinoma specimens prior platinum-based chemotherapy treatment, we demonstrate that SLFN11 is expressed by infiltrating innate and adaptive immune cells. We show, for the first time, that SLFN11 density in both the neoplastic and microenvironmental components was independently associated with favorable outcome. Transcriptomic analyses suggested the presence of a hitherto modulation of the cancer-immunity cycle orchestrated by SLFN11. We propose SLFN11 as a dual biomarker capturing simultaneously interconnected immunological and cancercell-intrinsic functional dispositions associated with sensitivity to DNA damaging agents.

## Introduction

The putative DNA/RNA helicase Schlafen11 (SLFN11) was independently reported by us(Zoppoli *et al*, 2012) and others to be the top correlating transcript, amongst more than 20,000, with the response of cancer cells to DNA damaging agents (DDA) with different modes of action such as topoisomerase I (e.g. topotecan and irinotecan)(Barretina *et al*, 2012; Coussy *et al*, 2020), topoisomerase II inhibitors (e.g. epirubicin) and bulk alkylating or alkylating and crosslinking-like agents (e.g. cyclophosphamide or platinum salts, respectively)(Conteduca *et al*, 2020; Iwasaki *et al*, 2019; Stewart *et al*, 2017). Subsequently, a positive association between SLFN11 and sensitivity to Poly (ADP-ribose) polymerase inhibitors (PARPi) was also described(Lok *et al*, 2017; Murai *et al*, 2016; Pietanza *et al*, 2018; Stewart *et al*., 2017). After our discovery, several studies confirmed the causal role of SLFN11 in the process of cell death upon DDA challenge in cell lines(Murai *et al*, 2018; Murai *et al*, 2020), organoids(Conteduca *et al*., 2020) and xenografts(Coussy *et al*., 2020; Iwasaki *et al*., 2019; Stewart *et al*., 2017) from different tumor types. Moreover, SLFN11 has been recently studied in relation with the immune system(Mezzadra *et al*, 2019; Stewart *et al*., 2017), especially in breast cancer(Isnaldi *et al*, 2019), and for its potential role as an endogenous inhibitor of viral replication(Li *et al*, 2012) and translation of DNA damage response proteins(Li *et al*, 2018). Taken together, the available literature suggests that SLFN11 may play a so far not completely understood role in an intertwined process of cancer and immune response to DDA-based chemotherapies. Indeed, it has been shown that SLFN11 is strictly correlated with immune-related transcripts in breast cancer(Isnaldi *et al*., 2019), and its expression is regulated by interferon signaling in primary human cells(Li *et al*., 2012; Puck *et al*, 2015) and, possibly, also in neoplastic cells(Mezzadra *et al*., 2019; Stewart *et al*., 2017). Moreover, SLFN11 is associated with early interferon-response genes in neoplastic cells(Stewart *et al*., 2017), hence pointing toward an exogenous regulation of its levels by the tumor-infiltrating immune milieu. One of the human cancers whose standard-of-care (SoC) treatment relies upon DDA, and which are considered particularly sensitive to such category of chemotherapeutics, is highgrade serous ovarian carcinoma (HGSOC). HGSOC is the most common histologic subtype of ovarian cancer, accounting for three quarters of newly diagnosed cases(Lisio *et al*, 2019). Initial SoC treatment for advanced stage HGSOC (the most frequent presentation stage for this poor-prognosis disease) consists of a platinum salt-taxane chemotherapy (CT) combination regimen, interposed or preceded by surgical debulking(Lheureux *et al*, 2019). In spite of macroscopically complete resection (R0) and upfront chemotherapy, most HGSOC patients will eventually progress and die from their disease. In this context, several studies have shown that tumor-infiltrating lymphocytes (TILs), especially CD3+ and CD8+ TILs, may have a role as a prognostic biomarker, but their clinical utility is still unclear(Stanske *et al*, 2018). In this study, our main aim was to determine whether SLFN11 transcript and protein could be accurately and reproducibly measured in two different serous ovarian cancer cohorts, one internal and another one from TCGA, considering the following aspects: a) the sensitivity of HGSOC to DDA, b) the need for clinically useful prognostic biomarkers for chemotherapy treatments, c) the potential connection between SLFN11 and TILs, and, d) the potential shown by SLFN11 modulation in preclinical models. We explored how SLFN11 protein is expressed in cancer cells and their surrounding microenvironment and, most importantly, whether SLFN11 could represent a relevant prognostic biomarker to platinum-based treatment response in advanced stage HGSOC patients.

## Results

### 1. Demographics

The clinico-pathological features of HGSOC cases selected for the present analysis, as detailed in the Methods section, are reported in Table 1 and Supplementary Table 1. The proportions of advanced stage HGSOC patients were balanced between platinum-resistant (PR, N = 13) patients, defined as progressing within six months from the end of first CT, and platinum-sensitive (PS, N = 15) ones (85% and 87% respectively), as was the median number of completed cycles (seven in both groups). Median progression-free interval (PFI) was 4 months (95% CI = 2 – 6) in PR patients and 11 months (95% CI = 9 – not reached) in PS ones. The patients had a median age of 62.4 years (95% CI = 56.8 – 66.9). In the studied cohort, PR cases were on average older than PS ones (65.7 vs. 58.2 years, p-value = 0.0026).

**Table 1:** Cohort demographics

|  | **PR** | **PS** | **p-value** |
| --- | --- | --- | --- |
| Stage (n, percent) |  |  |  |
| IIIc | 11 (85) | 13 (87) | 1.000 |
| IVa | 2 (15) | 2 (13) |  |
| Cycles (number, IQR) | 7 (6 – 8) | 7 (6 – 9) | 1.000 |
| PFI (months, 95% CI) | 4 (2 – 6) | 11 (9 – n.r.) | n.c. |
| Age at diagnosis (years, IQR) | 65.7 (64.3 – 69.8) | 58.2 (54.7 – 61.5) | 0.0026 |

PR = platinum-resistant, PS = Platinum-sensitive, IQR = interquartile range, PFI = progression-free interval, n.r. = not reached, n.c. = not calculated.

### 2. SLFN11 levels are precisely defined by both transcript and protein levels in HGSOC samples

To assess SLFN11 in HGSOC cases, we evaluated both the transcript levels by quantitative real-time polymerase chain reaction (qRT-PCR) as - ΔΔCt, and the protein levels by immunohistochemistry (IHC) as H-score, blindly measured by HALO (CW) in formalin-fixed, paraffin-embedded (FFPE) samples (see Supplementary Table 1 and Figure S1). Transcript and protein levels showed a strongly significant correlation (ρ = 0.52, p-value = 0.0051, see Figure 1A). This suggests that independent methods to measure SLFN11 in FFPE tissues yield comparable results, and that assessing either SLFN11 transcript or protein are both acceptable ways to analytically quantify the tissue levels of SLFN11 gene products. We next sought to test whether SLFN11 H-scores, blindly evaluated by a trained pathologist (JR) in cancer cells, could be consistently reproduced by digital pathology software such as HALO, which allows high content imaging assessment, and can provide quantitative measures on both cancer and non-cancer cells, as well as the two combined measures (“overall H-score”). Indeed, we found that the correlation between pathology-assessed and HALO-assessed H-scores in cancer cells was highly significant (ρ = 0.88, p-value < 0.0001, see Figure 1B), with excellent reliability(Koo & Li, 2016) (intraclass correlation coefficient – ICC – for agreement = 0.88 and ICC for consistency = 0.90, see Figure 1C), no relevant bias, and a very slight trend towards higher H-scores given by HALO for higher means, as evaluated with the Bland-Altman limits of agreement method (see Figure 1D). Taken together, these results established the analytical validity of our IHC approach to SLFN11 measurement in tumor specimens.

**Figure 1:**
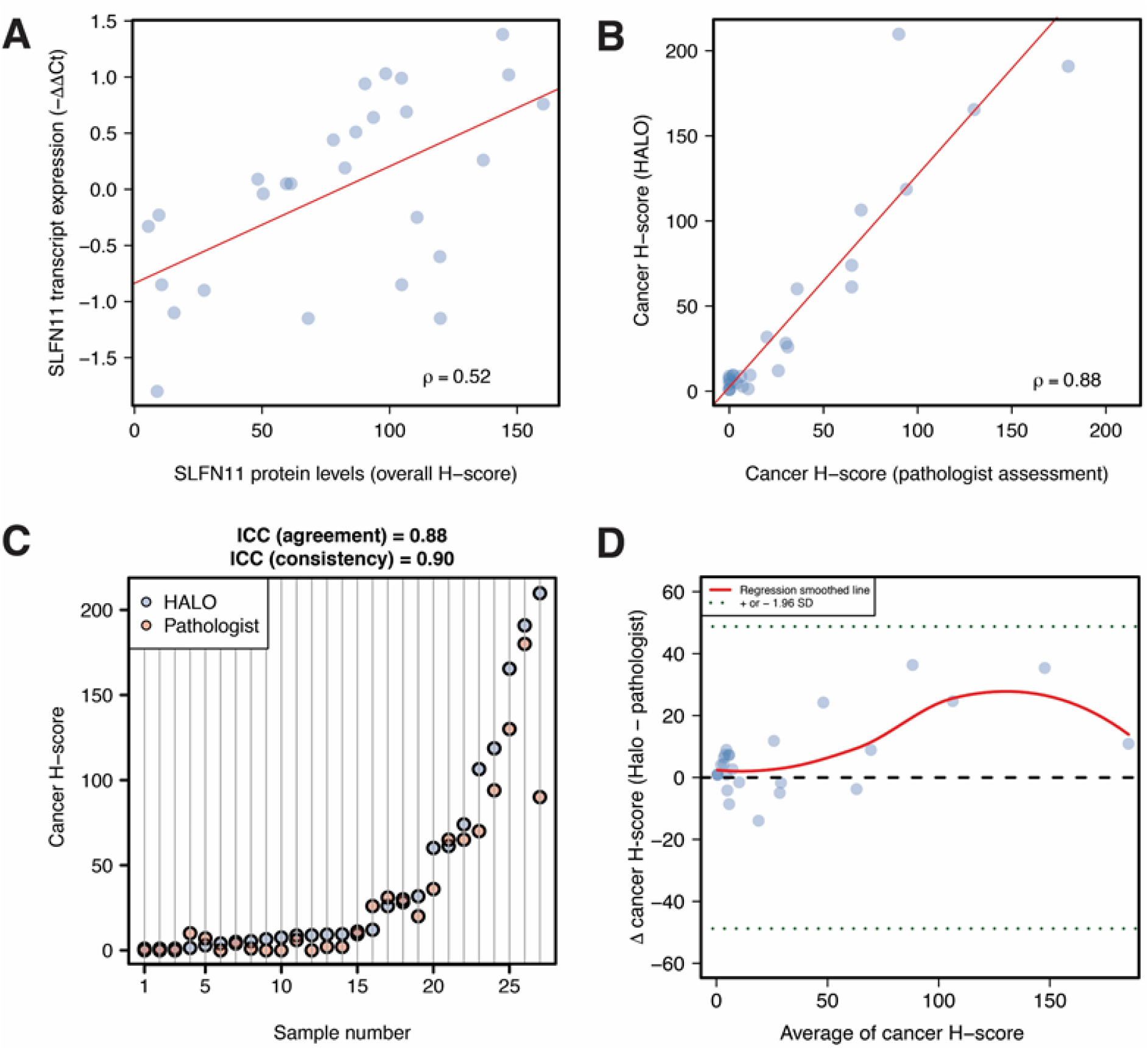
SLFN11 transcript and protein levels in HGSOC. ***Panel A:*** Scatterplot representing SLFN11 transcript by qRT-PCR as −ΔΔCt (*y* axis) as a function of its protein assessment by IHC as H-score (*x* axis) in non-cancer and cancer nuclear cells from HGSOC specimens; ρ is the Spearman’s correlation coefficient, the least squares regression is represented by the red line, whereas dots are measurements of SLFN11 by qRT-PCR and IHC in individual samples. ***Panel B:*** Scatterplot representing SLFN11 protein levels in HGSOC cancer cells. *X* axis: pathologist’s assessment; *y* axis: H-score measured by HALO Digital Pathology (DP) software. ***Panel C:*** Dot plot illustrating cancer-cell H-scores in individual samples (*y* axis), ordered by increasing DP-assigned values (*x* axis), highlighting the excellent consistency of intraclass correlation coefficients (ICC) between the two methods. Each dot represents a score assigned by either the DP software (HALO) or the pathologist performing the assessment. ***Panel D:*** Bland-Altman plot displaying the difference between HALO and pathologist’s H-scores for cancer cells (*y* axis) by the increasing mean of value couples for individual samples (*x* axis). All points lie within 1.96 standard deviations (SD - dotted green horizontal lines) from the mean difference (dashed horizontal black line), indicating no relevant bias between raters, and an insignificant trend toward higher H-scores for HALO as the mean values increase. The red line represents a smoothed regression (loess) fit of the actual mean scores.

### 3. Total and intratumoral infiltrating lymphocytes contribute to SLFN11 levels in HGSOC

Since several studies have reported on the role of TILs in the prognosis of ovarian cancer (Goode *et al*, 2017; Hwang *et al*, 2012; Li *et al*, 2017; Sato *et al*, 2005), and SLFN11 has been shown to be expressed in primary human T-lymphocytes(Puck *et al*., 2015), we evaluated TIL infiltration by CD3 and CD8 staining (analysis performed by FG), both in terms of total number/mm (total CD3+ and CD8+ TILs) and as a measure of TILs in direct contact with cancer cells, without stroma interposition (intratumoral CD3+ and CD8+ TILs) in HGSOC(Stanske *et al*., 2018). Using HALO, we then calculated SLFN11 overall H-scores in the studied cohort, as well as H-scores in cancer and non-cancer cells separately (see Supplementary Table 1). H-scores measured in non-cancer cells showed the strongest correlations with TILs, whereas the H-scores in cancer cells exhibited a non-significant negative association with TIL counts (see Figure 2A and Supplementary Table 2). In particular, non-cancer H-score correlations with total CD3+ and CD8+ TILs were moderate at 0.41 (false discovery rate - FDR = 0.0723, see Figure 2B) and 0.39 (FDR = 0.0852, see Figure 2C) respectively. Indeed, SLFN11 protein assessment in the clinical specimens revealed its localization in both cancer cells and stromal cells of various origins (see Figure 2D for representative images of stroma SLFN11-high and tumor SLFN11-high or SLFN11-low pictures). Taken together, these results indicate that, in addition to TILs, other cell populations contribute to SLFN11 protein levels in tumor tissues. Of interest, a moderate association between cancer and non-cancer SLFN11 levels could be observed (ρ = 0.50, FDR = 0.0208).

**Figure 2:**
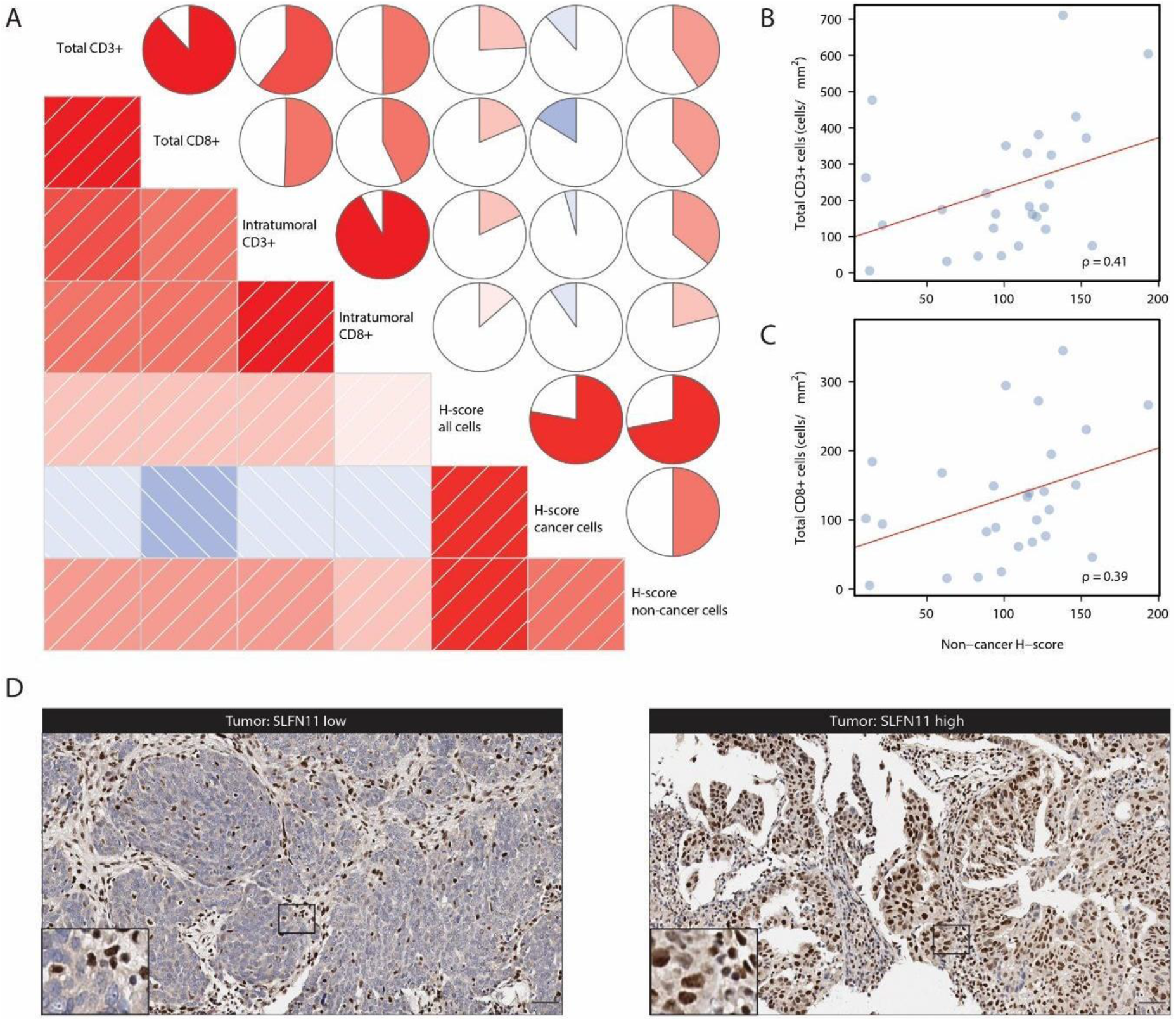
SLFN11 protein levels in HGSOC and their correlation with TILs. ***Panel A:*** Correlogram of TILs and SLFN11 H-scores, assessed in overall, cancer and non-cancer cells. In the lower triangle of the graph, boxes represent pairwise correlations colored by direction (blue for negative correlations and red for positive ones) and strength (intensity of shading) of the correlation itself. In the upper triangle, circles use the same scaled colors, but fill an area proportional to the absolute value of the correlation, and are filled clockwise for positive values, anti-clockwise for negative values. ***Panels B and C:*** Scatterplots representing total CD3+ cells – panel B – and total CD8+ cells – panel C – (*y* axes, cells/mm) as a function of SLFN11 H-score in non-cancer cells (*x* axis); ρ is the Spearman’s correlation coefficient, the least squares regression are represented by the red lines, whereas dots are measurements of immune cell counts by H-scores in individual samples. ***Panel D:*** Representative images of SLFN11 IHC in HGSOC specimens. Left, stroma SLFN11 high and tumor SLFN11 low and right, stroma and tumor SLFN11 high, for the indicated cancers. The insets highlight nuclear SLFN11 protein localization in tumor cells and different stromal cell subtypes. Scale bars, 50 μm. The insets show a 3x magnification of the representative image.

### 4. SLFN11 in cancer and non-cancer cells independently predicts response to platinum-based chemotherapy in HGSOC

We next sought to explore whether higher SLFN11 protein levels associate with better outcome in platinum-treated advanced stage in our HGSOC cohort. First, we evaluated the impact of SLFN11 overall H-score, H-score in cancer and non-cancer cells, as well as stage, age, and TIL infiltration on PFI by univariable statistics. Overall and non-cancer SLFN11 H-scores were strongly associated with a better prognosis (HR = 0.50, 95%CI = 0.33 – 0.75, p-value = 0.0009, and HR = 0.54, 95%CI = 0.36 – 0.81, p-value = 0.0028 respectively). Among the other variables with possible prognostic impact assessed in our cohort, older age was associated with shorter PFI (hazard ratio – HR = 1.83, 95% confidence interval – 95%CI = 1.19 – 2.82, p-value = 0.0062), whereas higher total CD3+ TILs were associated with longer PFI (HR = 0.55, 95%CI = 0.34 – 0.90, p-value = 0.0180). A borderline significant association with shorter PFI was observed for stage IVa vs. IIIc cancer (HR = 3.19, 95%CI = 0.89 – 11.46, p-value = 0.0758), whereas an opposite, trend be found for higher total CD8+ TIL count (HR = 0.67, 95%CI = 0.41 – 1.09, p-value = 0.1040) and SLFN11 H-score assessed in cancer cells only (HR = 0.62, 95%CI = 0.38 – 1.02, p-value = 0.0620). With an H-score cutoff of 60, obtained by maximizing the accuracy to classify PR versus PS cases in our cohort, overall SLFN11 protein levels had an accuracy = 0.78, with sensitivity = 0.93 and specificity = 0.62 (see Figure 3A). The association of overall SLFN11 H-score as a binary variable with PFI was indeed significant, with an HR = 0.17 (95%CI 0.06 – 0.45, p-value = 0.0004, see Figure 3B). When the most significant measure of SLFN11, i.e. the overall H-score, and the other variables with a p-value ≤ 0.1 were entered in a stepwise forward-backward multivariable Cox’s regression model, overall SLFN11 protein levels retained their independent prognostic value (adjusted HR = 0.56, 95%CI = 0.37 – 0.85, p-value = 0.0073), together with age and stage (see Figure 3C). Albeit exploratory in nature, these results were surprising in several regards. First, the independent prognostic value of overall SLFN11 H-score suggests that SLFN11 levels in both cancer and non-cancer cells may play a role in response to platinum-containing regimens in HGSOC. Indeed, dichotomized SLFN11 cancer (see Supplementary Figures 2A and 2B) and non-cancer (see Supplementary Figures 2C and 2D) levels were also prognostic by univariable analysis (even though with smaller significance than overall SLFN11, see Supplementary Table 3). Moreover, both cancer and non-cancer SLFN11 retained their independent role in stepwise multivariable models starting from the same set of variables as the one including overall SLFN11 (see Supplementary Figures 2E and 2F and Supplementary Table 3). An extremely interesting finding is that, when overall SLFN11 or non-cancer SLFN11 are considered together with the other covariates to generate a multivariable model, CD3+ TILs lost their independent prognostic role. On the other hand, when cancer SLFN11 H-score is used to generate a multivariable model, that measure is independently significant together with CD3+ TILs (see Supplementary Figure 2E). This observation strengthens the hypothesis that SLFN11 expressed in cancer cells, as observed in other studies, is directly linked with neoplastic cell sensitivity to alkylating agents independently of immune infiltration, whereas the prognostic relevance of non-cancer SLFN11 may instead underlie an “active” tumor immune milieu. In turn, these findings would explain why overall SLFN11 H-score is a stronger prognostic biomarker than either cancer or non-cancer SLFN11 measured separately.

**Figure 3:**
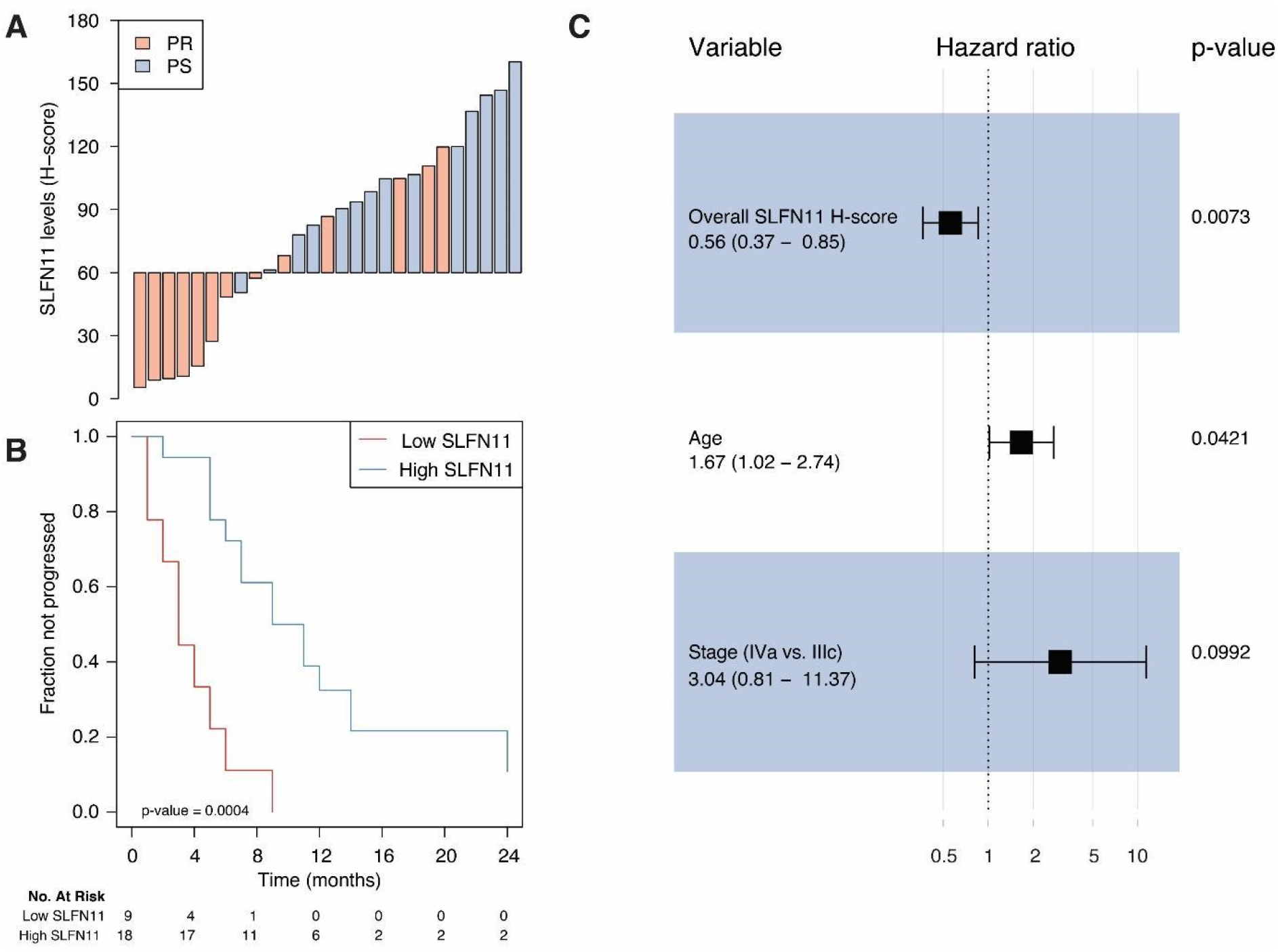
SLFN11 protein levels measured in cancer and non-cancer cells are independently prognostic in HGSOC. ***Panel A:*** Waterfall plot showing SLFN11 overall protein levels (i.e., measured in cancer and non-cancer cells) in individual cases, colored by platinum sensitivity: SLFN11 protein is reported as H-score (y axis), whereas cases are reported by increasing values (x axis) and colored in red if platinum-refractory (PR) or light blue if non-refractory (NR). ***Panel B:*** Kaplan-Meier plot showing progression-free interval (PFI) stratified by SLFN11 overall protein levels (“high” if H-score > 60, “low” if < 60). The progressed fraction of patients (*y* axis) is plotted against time expressed in months from the end of first-line chemotherapy, censored at 24 months (*x* axis). Numbers at risk are reported below the plot. P-value in the bottom left of the plot is from the Wald statistics for the univariable Cox’s regression. ***Panel C:*** Forest plot of hazard ratios (x axis, in log scale) for variables retained in the final multiple Cox’s regression model. Point HR estimates are reported below each variable together with 95% confidence intervals (95%CI) in parentheses, whereas adjusted p-values for each variable are on the right side of the plot. Filled black squares represent HR estimates, with relative 95%CI shown as horizontal lines with brackets.

### 5. SLFN11 is expressed by cells of the innate and adaptive immune system infiltrating HGSOC

As our results indicated, SLFN11 plays a role in the response to platinum-based treatment in HGSOC, due to its expression in both cancer and non-cancer (immune-related) cells. Moreover, non-cancer SLFN11 H-score shows only a moderate correlation with TILs, and they retain an independent prognostic value in HGSOC, thus suggesting that other cell subpopulations contribute to the overall levels of SLFN11 in tissues. Hence, we sought to better define these populations. With this aim, we estimated cancer cellularity in a second HGSOC cohort from The Cancer Genome Atlas (TCGA N = 302 cases) with ESTIMATE(Yoshihara *et al*, 2013), and we inferred leukocyte subpopulations using CIBERSORTx, a well-established method for characterizing the immune cell composition of tissues from their gene expression profiles(Newman *et al*, 2015). We first correlated the obtained values with SLFN11 transcript levels (see Supplementary Table 4). Of interest, not only adaptive immune system cells (CD4+, CD8+ T-cells as well as B cells), but also macrophages and Natural Killer (NK) cells showed a significant association with SLFN11 (FDR < 0.05). In fact, amongst immune cell subpopulations, macrophages showed the strictest correlations with SLFN11, and such observation was transversally confirmed for validation purposes by single sample gene set enrichment analysis (ssGSEA, correlation between enrichment score for macrophages and SLFN11 = 0.27, p-value < 0.0001, see Supplementary Figure 3A). Moreover, publicly available RNA-sequencing data from sorted leukocyte subpopulations (GEO accession GSE60420) corroborated our *in silico* findings: higher SLFN11 levels were observed in monocytes, followed by NK cells, CD8+ T-cells, B-cells and CD4+ T-cells, whereas - as in our results - the lowest SLFN11 transcript could be observed in neutrophils (see Supplementary Figure 3B). Cancer cellularity showed a negative correlation with SLFN11 expression in HGSOC TCGA OVCAR (ρ = −0.30, FDR < 0.0001, see Supplementary Figure 4A). This observation, independently confirmed in our cohort (ρ = −0.42, p-value = 0.0276, see Supplementary Figure 4B), could be explained by hypothesizing that HGSOC cases with lower cancer cellularity would, in general, have higher tumor immune cell infiltration, with consequently higher SLFN11 levels. To explore this hypothesis, we performed principal component analysis (PCA), including cancer cellularity together with immune cell subpopulations correlating with SLFN11 with FDR < 0.05 (see Figure 4A). PCA is a dimensionality reduction method enabling the identification and visualization of correlations and patterns between variables without aprioristic assumptions about their mutual relationships. As anticipated, all immune subpopulations are represented as lying in opposition to cancer cellularity, in particular M1 macrophages, CD4+ memory resting T-cells and CD8+ T-cells. Moreover, the SLFN11 expression vector is lying close to those of the immune subpopulations it is more closely correlated to, such as T-cells and macrophages. Overall, PCA substantiates our hypothesis that high immune infiltration is driving the negative correlation of SLFN11 with cancer cellularity and that specific immune subpopulations are closely associated with high SLFN11 expression in HGSOC.

**Figure 4:**
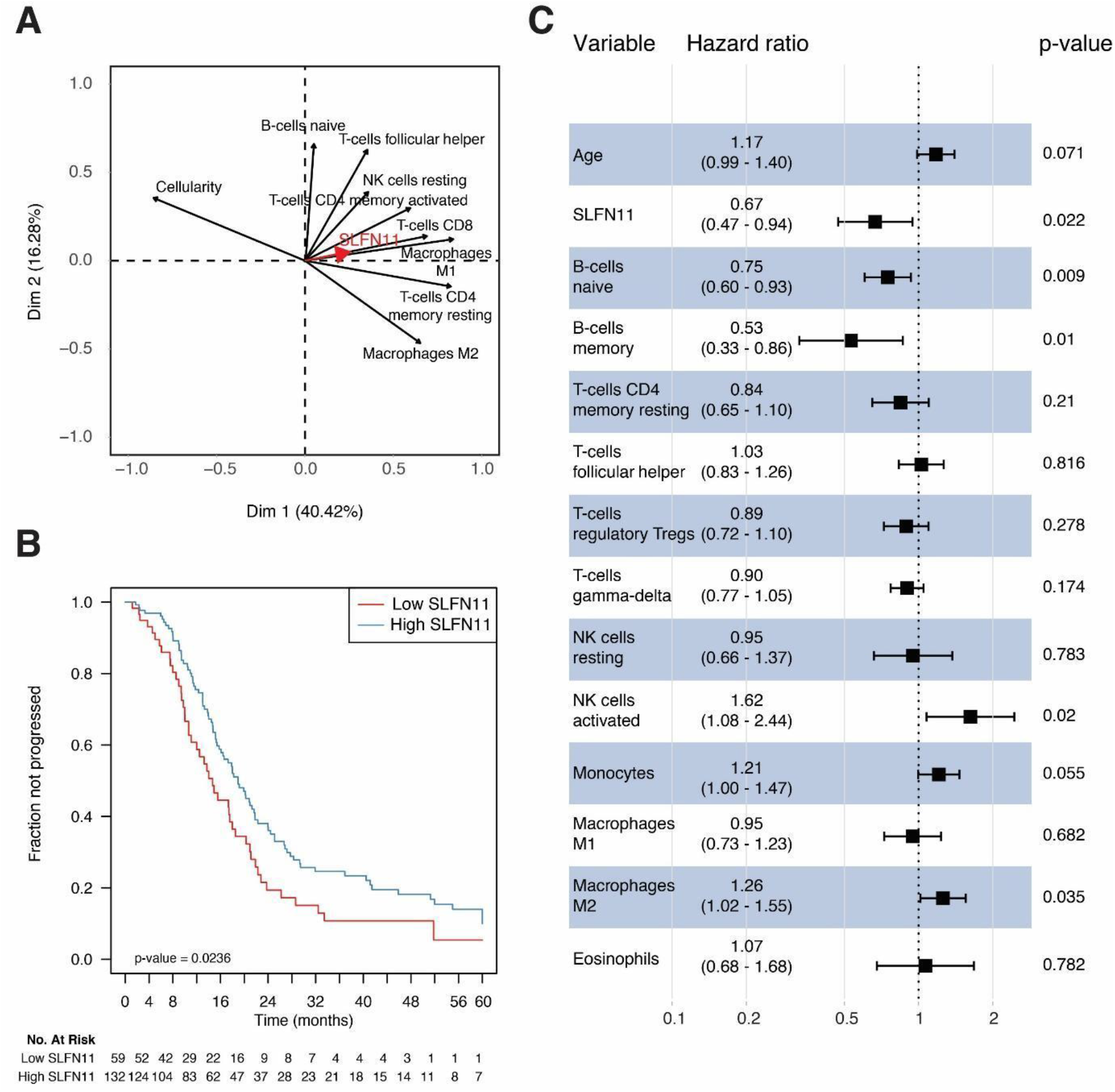
SLFN11, immune cell subpopulations and prognosis in the TCGA serous ovarian carcinoma dataset. ***Panel A:*** Variable correlation plot of the principal component analysis including SLFN11 transcript, cancer cellularity, and CIBERSORTx immune cell subpopulations significant by univariable correlation at FDR < 0.05. The two axes represent the first principal components explaining the greatest fraction of the variance of the analysed dataset, with percent of explained variability in parentheses. The relative position of the variables towards each other explains their relative correlation, whereas their distance from the intersect accounts for their contribution to the components. SLFN11 is represented with a thick red arrow for sake of clarity. ***Panel B:*** Kaplan-Meier plot showing progression-free interval (PFI) stratified by SLFN11 transcript (“high” if above the lower tertile of expression in the dataset, “low” if below). The progressed fraction of patients (*y* axis) is plotted against time expressed in months from the end of first-line chemotherapy, censored at 60 months (*x* axis). Numbers at risk are reported below the plot. P-value in the bottom left of the plot is from the Wald statistics for the univariable Cox’s regression. ***Panel C:*** Forest plot of hazard ratios (*x* axis, in log scale) for variables retained in the lasso-selected multiple Cox’s regression model. Point HR estimates are reported below each variable together with 95% confidence intervals (95%CI) in parentheses, whereas adjusted p-values for each variable are on the right side of the plot. Filled black squares represent HR estimates, with relative 95%CI shown as horizontal lines with brackets.

### 6. SLFN11 is independently prognostic in the TCGA HGSOC data set

Finally, we validated the prognostic role of SLFN11 in TCGA HGSOC patients. To do so, we selected stage IIIc/IV cases with histological grade 3 and at least 28 days of progression-free interval (PFI - 221 cases with 157 progression events). SLFN11 was associated with PFI (HR = 0.68, 95%CI = 0.49 – 0.95, p-value = 0.0233, see Figure 4B), and remained independently significant (adjusted HR = 0.67, 95%CI = 0.47 – 0.94, p-value = 0.0222), together with age and specific immune cell subpopulations, in a multivariable Cox’s proportional hazards model with variables selected by lasso regularization (see Figure 4C). Of relevance, B and T-cell subpopulations were independently prognostic of an extended PFI, whereas monocytes, M2 (but not M1) macrophages, and activated NK cells were associated with poorer prognosis (for univariable survival analyses of immune cell subpopulation in the TCGA data set, see Supplementary Table 5). This result is in line with SLFN11 correlations with immune cell subpopulations, in that B-cells and NK resting cells were associated with higher SLFN11 transcript, while NK activated cells were not. The surprisingly negative association between NK and prognosis might be related to imbalance of different NK cell subsets associated with diverse immune-modulatory properties. Monocytes and macrophages exhibited heterogeneous behavior in regard with prognosis when assessed in a multivariable fashion. We did not, however, try to model the interactions of SLFN11 with those subpopulations, due to the interpretational complexity of results in the absence of functional experiments, as well as the limited numerosity of the TCGA HGSOC dataset for such a purpose. Likewise, we wished to avoid the overinterpretation of an exploratory analysis which would, anyway, be derived from *in silico* deconvolution methods not devoid of the potential for error propagation.

### 7. SLFN11 protein localization with immune cell subpopulations is confirmed in tonsil and HGSOC tissues

SLFN11 protein localization in a subset of cells in the immune cell compartment could be confirmed in tonsil and HGSOC (Figure 5) tissues from our cohort. In the tonsil, SLFN11 positive cells could be mainly found in the germinal center and the paracortical zone, and less in the mantle zone, of the lymphoid follicle. The germinal center is enriched in naïve and memory B-cells (CD20+) and monocyte/macrophages (CD68+), whereas the paracortical zone is mostly composed of T-cells (e.g. CD3+ and CD8+) (Figure 5A and supplementary figure 5). In HGSOC tissues we could observe high SLFN11 protein in those immune cell subtypes, particularly in monocytes/macrophages (Figure 5B, C for representative HGSOCs with varied SLFN11 protein in cancer cells). However, we also noted SLFN11 in other, to be defined, stromal cell subtypes. Taken together, we confirm that SLFN11 is expressed in macrophages, TILS and B-cells, but also in other still to be defined stromal cell types in HGSOC.

**Figure 5:**
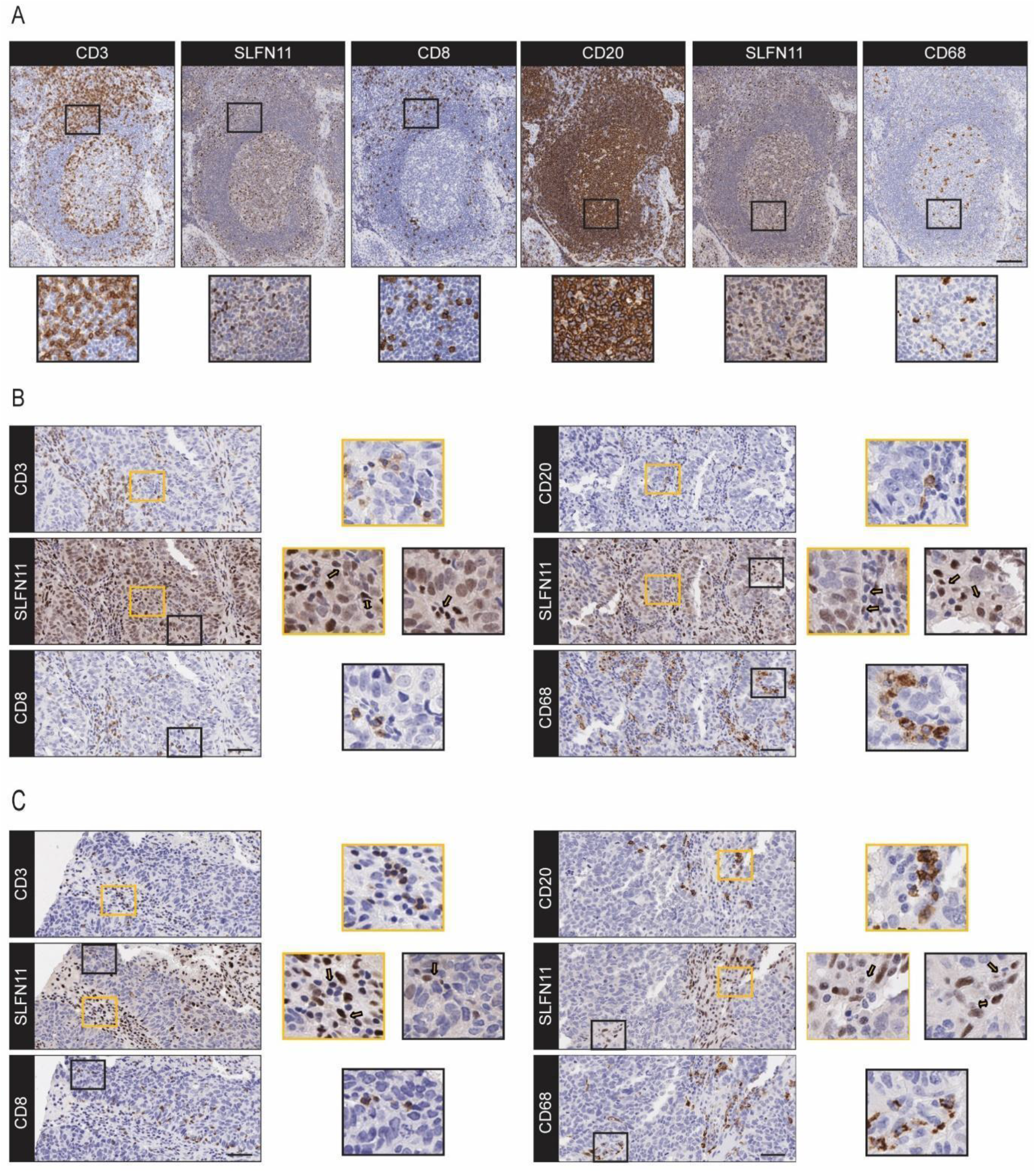
SLFN11 is expressed in a subset of immune-related cells in tonsil and HGSOC tissues. ***Panel A:*** Representative images of CD3, CD8, SLFN11, CD20 and CD68 IHC on serial sections of tonsil tissue. Shown is a lymphoid follicle with the round-to oval shaped germinal center, the surrounding mantle zone and, at the outer layer of the lymphoid follicle, the paracortical zone. SLFN11 is mainly expressed in the germinal center, which is mostly composed of B-cells (CD20+) and macrophages/monocytes (CD68+), as well as in the T-cell rich (CD3+/CD8+) paracortical zone. ***Panels B and C:*** Representative images of CD3, CD8, SLFN11, CD20 and CD68 IHC on serial sections of tumor SLFN11 high- (Panel B) and low (Panel C) and stroma SLFN11 high cancers. The insets show nuclear SLFN11 in cytosolic/membrane-based CD3, CD8, CD20 and CD68 positive cells (indicated by arrows in Panel B and C). Scale bars, 100 μm (Panel A) and 50 μm (Panel B). The insets show a 3x magnification of the representative images.

### 8. High SLFN11 expression is associated with immune activity signatures in HGSOC

After demonstrating, *in silico* and through immunohistochemical analysis on an independent case cohort, that SLFN11 is not only expressed in cancer cells, but also in TILs, in macrophages, and other immune cell subpopulations, we wondered how SLFN11 expression in HGSOC is correlated with biologically selected, well-established immune signatures representing hallmarks of immune activity, such as interferon α and γ signaling and STAT1 activation(Thorsson *et al*, 2018) (Liberzon *et al*, 2015; Teschendorff *et al*, 2010), MHC I and MHC II upregulation(Rody *et al*, 2009; Rody *et al*, 2011), antigen presenting machinery(Senbabaoglu *et al*, 2016), immune xonstant of rejection(Bedognetti *et al*, 2016; Hendrickx *et al*, 2017) and immunogenic cell death(Garg *et al*, 2016). To do so, we correlated SLFN11 with those signatures in the HGSOC TCGA RNA-seq dataset (N=302 cases). All the aforementioned signatures showed extremely significant positive correlations with SLFN11 expression, with close reproducibility among signatures of similar meaning generated by different authors (see Figure 6). In particular, the two top correlating signatures with SLFN11 in HGSOC were the immunogenic cell death signature (FDR = 5.44 x 10) and the interferon γ response hallmark signature (FDR = 1.23 x 10). Taken together, these findings are suggestive of a close, hitherto scarcely investigated link between SLFN11 and cancer immunity in HGSOC.

**Figure 6:**
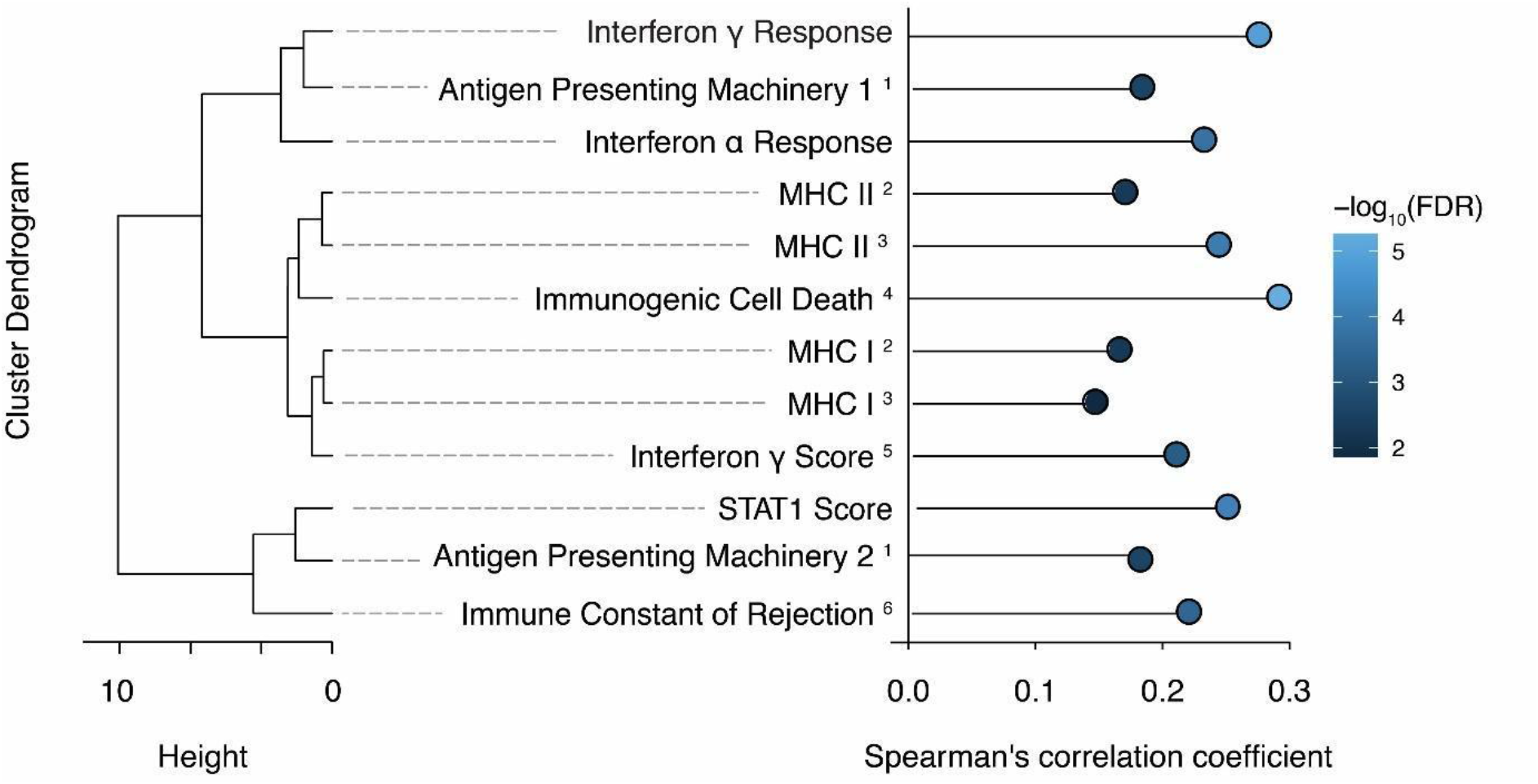
SLFN11 expression is associated with immune signatures in HGSOC. ***Left:*** Dendrogram representing the similarity between different immunologic signatures calculated in the TCGA ovarian cancer dataset (N = 302 cases). *x* axis represents the Ward’s D2 distance. Signature names have superscript numbers to denote the publication they are derived from: 1: Senbabaoglu Y et al., 2016(Senbabaoglu *et al*., 2016); 2: Rody A et al., 2009(Rody *et al*., 2009); 3: Rody A et al., 2011(Rody *et al*., 2011); Garg AD et al., 2016(Garg *et al*., 2016); Teschendorff AE et al., 2010(Teschendorff *et al*., 2010); if not specified, signatures are from 6: Thorsson V et al., 2018(Thorsson *et al*., 2018). ***Right:*** lollipop plot of correlations between SLFN11 and the aforementioned signatures. *x* axis represents the Spearman’s correlation coefficient between SLFN11 expression in the TCGA ovarian cancer dataset and the investigated signatures, whereas individual dots are coloured by −log10 of the FDR of the correlation.

## Discussion

Since the discovery of the human *SLFN* isoform SLFN11 in 2009(Bustos *et al*, 2009), SLFN11 has been reported to play a role in the native immune response such as viral infections and interferon(Li *et al*., 2012; Puck *et al*., 2015), as well as a potential role in adaptive immunity in cancer(Isnaldi *et al*., 2019; Stewart *et al*., 2017). In addition, work from various groups confirmed SLFN11 as being a determinant of sensitivity to a broad range of DDAs with different modes of action(Barretina *et al*., 2012; Conteduca *et al*., 2020; Coussy *et al*., 2020; Deng *et al*, 2015; Iwasaki *et al*., 2019; Stewart *et al*., 2017; Zoppoli *et al*., 2012), as well as PARP inhibitors(Lok *et al*., 2017; Murai *et al*., 2016; Pietanza *et al*., 2018; Stewart *et al*., 2017), in different, mainly preclinical, cancer settings. An understanding how SLFN11 modulates the response to chemotherapy in patients is of paramount importance for both basic biology and clinical viewpoints and is currently missing.

The assessment of SLFN11 as a clinical biomarker is hindered by the lack of validate algorithms to score it in human tissues. By implementing analytic pipelines in clinical material, we demonstrate that SLFN11 in both the neoplastic and microenvironmental compartments of tumor specimens modulates the response to platinum-containing regimens in HGSOC patients. Our proposed working model, based by integrating our data with current knowledge, is as follows (Fig. 7): SLFN11 expression might be controlled by endogenous (i.e. methylation) and exogenous factors (i.e., interferon gamma produced by tumor-reactive T cells). SLFN11 high cancer cells are more prone to undergo immunogenic cell death spontaneously or following platinum or other DNA damaging agent’s administration(Garg *et al*, 2017). Accordingly, T-cells are recruited and activate the interferon signaling cascade with consequent upregulation of MHC I and MHC II molecules, and induction of the cancer-immunity cycle. SLFN11 in cancer cells may directly contribute to immune system activation by inducing tumor necrosis factor- and innate immune response pathways(Murai *et al*., 2020) and interferon signaling may directly and indirectly prompt cytotoxicity in cancer cells(Mezzadra *et al*., 2019).

**Figure 7:**
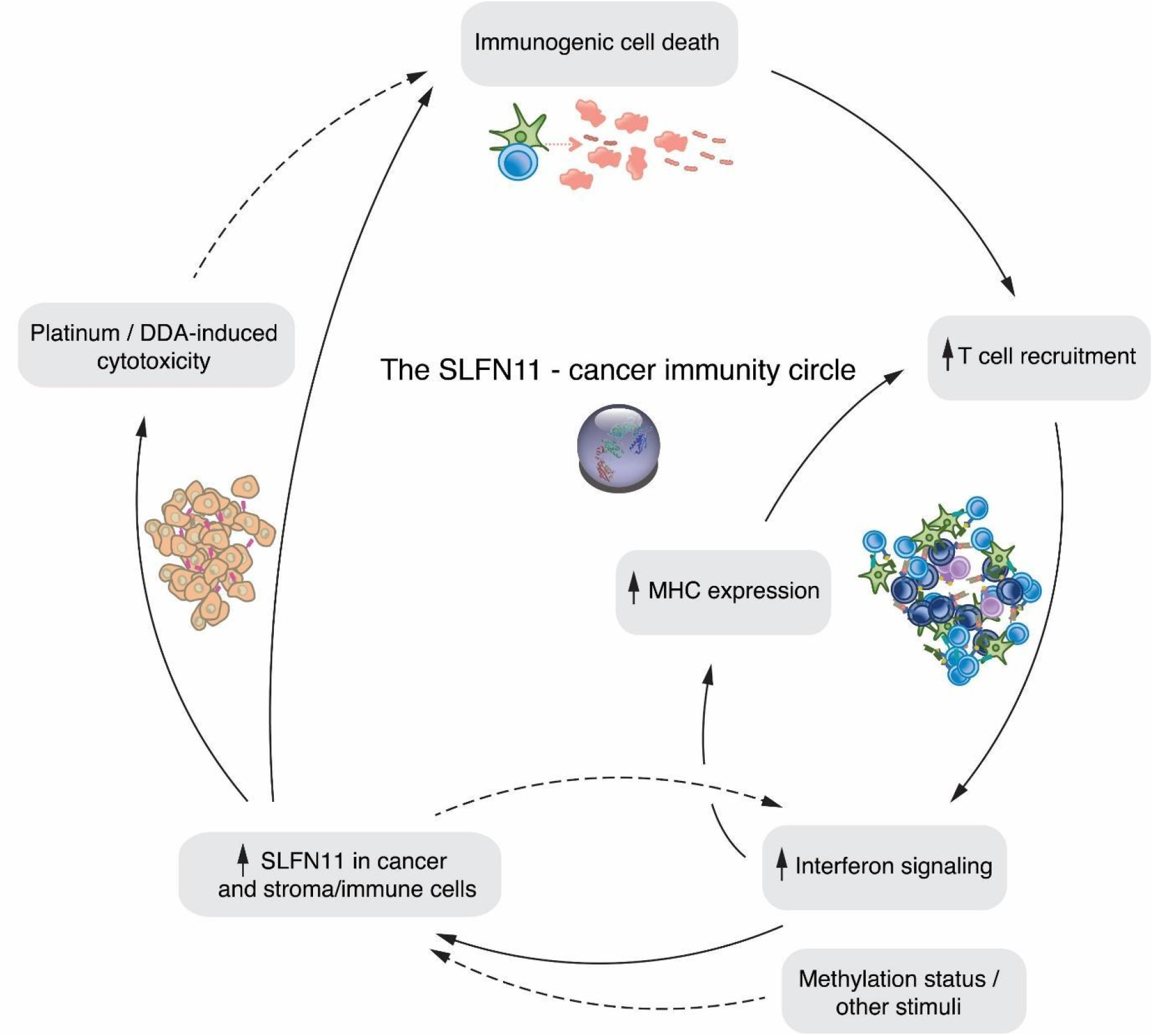
Proposed model for the SLFN11-cancer immunity cycle. Clockwise from bottom left: SLFN11 can be upregulated in cancer cells and tumor infiltrating immune cells by interferon signaling and possibly by other factors, such as SLFN11 promoter demethylation. SLFN11 high cancer cells are more prone to undergo immunogenic cell death spontaneously or following platinum or other DNA damaging agent’s treatment(Garg *et al*., 2017). This, in turn, recruits T cells, which activate the interferon signaling cascade and consequently stimulate the upregulation of the MHC I and MHC II complexes, and induction of the cancer-immunity cycle. Increased SLFN11 in cancer cells may directly contribute to immune system activation(Murai *et al*., 2020) and interferon signaling may directly and indirectly induce cytoxicity in cancer cells(Mezzadra *et al*., 2019). Dashed arrows indicate relations that were not assessed in the present manuscript but derived from other publications.

SLFN11 has been shown to predict response to DDA in different cancer models(Conteduca *et al*., 2020; Coussy *et al*., 2020; Deng *et al*., 2015; Iwasaki *et al*., 2019; Shee *et al*, 2019; Zoppoli *et al*., 2012), but to our knowledge we describe here for the first time that SLFN11 has more predictive power when its expression in the stromal non-cancer compartment is taken into account. These results are important in several regards. First, they support overall SLFN11 profiling in clinical tissues, and we provide evidence that levels of SLFN11 can be precisely measured by both transcript and protein analyses. Second, these findings point out that SLFN11, which is only expressed in humans and some primates, should be best assessed in clinical tissues, rather than xenograft- or *in vitro* cancer tissues, to understand its relevance in a clinical setting.

For the first time we confirm SLFN11 expression in tumor infiltrating innate (macrophages and NK cells) and adaptive immune cells (CD4+, CD3+ and CD8+ lymphocytes and B-cells). Our results support and add to recent findings, where it has been proposed that SLFN11 plays a role not only in the innate immune response such as defense mechanisms against viruses(Li *et al*., 2012) or damaged DNA(Mu *et al*, 2016; Murai *et al*., 2018; Murai *et al*., 2020), but also in adaptive immunity to combat cancer(Isnaldi *et al*., 2019; Stewart *et al*., 2017). Accordingly, Isnaldi et al. (2019) described an association of SLFN11 with TILs in breast cancer. Our results indicate high SLFN11 expression in macrophages and monocytes, in line with a previous report(Puck *et al*., 2015), whereas low expression was observed in neutrophils. SLFN11 protein localization was confirmed in macrophages, CD3+ and CD8+ TILs and naïve and memory B-cells in HGSOC tissues. Macrophages, when polarized to a M1 phenotype, are important to sustain a cytotoxic T cell response(Sica & Mantovani, 2012). However, we also noted SLFN11 presence in other stromal cell types that remain to be identified. Moreover, as SLFN11 is expressed in several infiltrating immune cells, it holds promise as a potential biomarker for immune infiltration and for response to immunotherapy drugs, although this warrants further investigations.

In our analyses, we demonstrate that SLFN11 in cancer and immune cells is independently a predictor of response to chemotherapy in HGSOC. These observations are in agreement with recent findings. Accordingly, in two other studies SLFN11 has been shown to be a predictive biomarker of longer survival in chemotherapy-treated ovarian cancer patient cohorts(Shee *et al*., 2019; Zoppoli *et al*., 2012). The concept that the immune-infiltrating milieu can modulate cancer prognosis is not new: multiple studies have reported on the role of TILs in the prognosis of ovarian cancer and HGSOC(Goode *et al*., 2017; Hwang *et al*., 2012; Li *et al*., 2017; Sato *et al*., 2005) and Liu et al. (2020) expanded this notion by looking at other leukocyte subpopulations(Liu *et al*, 2020). In analogy with these studies, we found that SLFN11 and TILs were associated with a better prognosis in our HGSOC cohorts. Similar trends were noted for naïve B-cells and macrophages of type 1; on the contrary, macrophages of type 2 were associated with poorer prognosis. We also noted that SLFN11 is correlated with well established, biologically selected, immune signatures (Fig. 6) in HGSOC. All together, these observations hint at the potential presence of interactions between SLFN11 and the immune system in cancer.

Potential interactions between SLFN11 and the immune system were proposed in literature. For example, SLFN11 has been described to be an IFN-stimulated gene in human blood mono-nuclear cells (PBMC)(Li *et al*., 2012) and primary human immune cells (monocytes and monocyte-derived dendritic cells)(Puck *et al*., 2015), and possibly also in cancer cells, fostering further investigations. Increased cancer SLFN11 (modulated by other factors like promoter demethylation(Nogales *et al*, 2015)), has been proposed to activate immediate early genes to stimulate interferon signaling in response to replication stress and DNA damage(Murai *et al*., 2020). In a different study, SLFN11 was identified as a sensitizer of tumor cells to T cell- and IFN-γ-mediated cytotoxicity.(Mezzadra *et al*., 2019) Accordingly, following IFN-γ exposure SLFN11 has been shown to couple interferon-γ receptor (IFNGR) signaling to the induction of DNA damage and cell death in tumor cells in a context-dependent fashion. Taken together, these results point at a complex interplay between SLFN11 in cancer cells and the immune system in cancer, which merits enticing further investigation.

We are aware of the limitations of our study. Among them, the retrospective nature of the analysis is unavoidable. The second limitation is the small sample size of our clinical cohort. Nevertheless, the fact that our observations are translated to a larger HGSOC cohort from TCGA, is assuring. Thirdly, even though SLFN11 expression in different leukocyte subpopulations could be confirmed in serial sections of tonsil and HGSOC tissues, these observations remain to be further validated by a multiplexing IHC approach. Nonetheless, to our knowledge, we show here for the first time in an *in silico* effort and in direct measurement SLFN11 presence and localization in tumor- and the tumor infiltrating milieu, in HGSOC patients.

In summary, the current study shows that SLFN11 in both cancer cells and a multitude of immune cells and potentially other (to be defined) cell types is associated with a better survival of HGSOC patients treated with platinum-containing regimens. Our findings add important information on the action of SLFN11 beyond its recently described role hence, we propose SLFN11 as a dual biomarker capturing simultaneously interconnected immunological and cancer-cell-intrinsic functional dispositions associated with sensitivity to DNA damaging agents.

## Patients and methods

### Study design

The objective of this study was to identify the association of SLFN11 with immune infiltration and prognosis in advanced high-grade serous ovarian carcinoma (HGSOC) patients undergoing neoadjuvant chemotherapy containing platinum. SLFN11 transcript was evaluated by RT-PCR and SLFN11 protein and immune infiltration localization by IHC. SLFN11 transcript and protein from noncancer and cancer cells were correlated with progression-free interval (PFI, defined as the time elapsing from the end of first treatment to clinical and/or radiological progression) to assess the value of SLFN11 as a biomarker of response to first-line platinum-containing chemotherapy regimens in HGSOC. Results were validated with a larger HGSOC cohort from TCGA.

### Patients

Patients receiving a diagnosis of HGSOC, treated with neoadjuvant platinum-based chemotherapy at Léon Bérard Cancer Center, Lyon FR, from January 2008 to June 2014, and meeting the following criteria, were retrospectively included, in a consecutive fashion, for the reported analyses: written informed consent for biobanking and use of samples for research purposes according to the Hosting Institution, histologically confirmed HGSOC (grade 3 according to the AJCC TNM stage(Amin *et al*, 2017), radiological and/or surgical classification as stage IIIc/IVa (FIGO classification)(Berek *et al*, 2018) at diagnosis, Eastern Cooperative Group (ECOG) performance status 0-1 at diagnosis, postmenopausal status at diagnosis, platinum-based neoadjuvant treatment at diagnosis followed by surgery, availability of a formalin-fixed, paraffin-embedded (FFPE) pre-chemotherapy tumor block from diagnostic biopsy, availability of clinical information concerning treatment and response duration and disease status at the time of sample collection. Cases were excluded: if they had primary debulking surgery followed by chemotherapy, if they were stage IVb, if they had received a previous or concomitant diagnosis of neoplasia (with the exclusion of carcinoma in situ of the cervix or skin basalioma) or if patients had received previous chemotherapies for any reason. Patients were divided into platinum-sensitive and platinum-resistant groups as previously described(Friedlander *et al*, 2011; Wilson *et al*, 2017). The presented research was conducted according to the ethical considerations and in compliance with the principles of the Declaration of Helsinki, approved by Regione Liguria Ethics Committee with registration number 347/2018 (approved 19/06/2019).

### RT-PCR

SLFN11 transcript quantification by q-RT-PCR was performed as previously described(Garutia *et al*, 2014). In brief, three membrane glass slides (PEN Membrane Glass slides, Arcturus^®^ Bioscience Inc. CA, USA), loaded with 8-μm thick sections were cut from FFPE-embedded pre-treatment diagnostic biopsies. One section of each sample was hematoxylin/eosin-stained and examined to assess tumor cellularity. Samples with a < 70% cellularity were subjected to microdissection. Upon RNA extraction and retro-transcription, RNA was quantified by Q-RT-PCR using a SLFN11-specific InvitrogenTM TaqMan^®^ assay (Invitrogen Inc. CA, USA) on an Applied Biosystems Inc. HT-7900 instrument. Samples were analysed in triplicate, using RPLP0, GAPDH and GUS as housekeeping (HK) genes. The mean PCR cycle thresholds (Ct) of the three HK genes was subtracted from the Ct value of SLFN11 for each sample, expressed as log_2_ (ΔCt) and, in turn, the median of ΔCts from the dataset was subtracted from ΔCts of single samples and inverted, to obtain a normally distributed, zero-centered semi-quantitative value for each sample (−ΔΔCt), as previously described(Barretina *et al*., 2012).

### RNA-sequencing profiling of human leukocyte subpopulations

RNA-sequencing results for SLFN11 in sorted leukocyte subpopulations from patients with immune-associated diseases, as further described by Linsley PS et al(Linsley *et al*, 2014), were obtained from the Gene Expression Omnibus (GEO, accession number GSE60424).

### Immunohistochemistry (IHC) and image analysis for SLFN11, CD3, CD8, CD20 and CD68

SLFN11 IHC was performed on 4 μM thick sections of formalin fixed paraffin embedded (FFPE) tissues and carried out on Bond RX (Leica Microsystem) using ER1 (pH6, Leica) antigen retrieval. Slides were stained with primary rabbit polyclonal anti-SLFN11 antibody (Abcam ab121731) at 2.5 μg/ml. Detection was performed with anti-rabbit poly-HRP-IGG, DAB refine and DAB enhancer (Leica, polymer refine detection kit, Leica). Digital slide images were acquired with the Aperio AT2 scanner (Leica) using a 20x objective. A HALO (Indica Labs) cytonuclear image analysis algorithm was optimized and run alongside different tissue classifiers and annotations, to capture the percentage of cancer, non-cancer and overall (cancer + non-cancer) nuclei with strong (3+), moderate (2+), weak (1+) or negative staining to calculate SLFN11 H-scores as [(%1+ cells) + (%2+ cells * 2) + (%3+ cells * 3)] in each sample. Samples were H&E stained to identify cancer cells. The same algorithm was used across all specimens and the analysis was blindly performed. All samples were in addition manually evaluated by a pathologist for cancer SLFN11 H-scores. CD3 and CD8 IHC was performed on full thick sections of FFPE tissues and carried out on Ventana Benchmark Ultra (Ventana Medical Systems) using heat-based antigen retrieval. Slides were stained with primary rabbit monoclonal anti-CD3 antibody (clone 2GV6 at 2.5 μg/ml) and primary rabbit monoclonal anti-CD8 antibody (clone SP57 at 2.0 μg/ml), both from Ventana Medical Systems. Slides were evaluated by a pathologist for total and intratumoral CD3 and CD8 by calculating mean CD3 and CD8 values from three high power field regions per sample. To confirm spatial resolution of SLFN11 protein in immune infiltrating cells, 4μm serial FFPE sections of human tonsil tissue or HGSOC specimens were taken. Sections were IHC stained for CD3, isotope control, SLFN11, CD8, CD20 and CD68 with BOND RX using ER1 (CD8 and CD20) or ER2 (CD3, IGG, SLFN11 and CD68) antigen retrieval. Primary antibodies used were as following: anti-SLFN11 as described above, anti-isotype control (ab172730, Abcam, at 2.5 μg/ml), anti-CD3 (clone 2GV6, Roche, at 0.4 μg/ml), anti-CD8 (clone C8/144 B, Dako, at 157 μg/ml), anti-CD20 (clone L26, Abcam, at 33.3 μg/ml) and anti-CD68 antibodies (clone PG-M1, Dako, at 0.3 μg/ml). Detection was performed with poly-HRP-IGG, DAB refine and DAB enhancer (polymer refine detection kit, Leica). Digital slide images were acquired with the Aperio AT2 scanner (Leica) using a 20x or 40x objective.

### Statistical analyses

Correlations between continuous variables were calculated using the Spearman’s rank coefficient and represented using scatter plots (package CNtu(Desmedt *et al*, 2016)), whereas differences in continuous distributions were calculated using the Wilcoxon test without continuity correction. Intraclass correlation coefficients (ICCs) were calculated to assess the consistency and agreement of IHC assessments (package psy(Shrout & Fleiss, 1979)), and visually inspected for bias and trend using dot plots and Bland-Altman plots. Correlation matrices and correlograms of IHC and TILs were generated using the package corrgram(Friendly, 2002), and multiple tests for association were adjusted using the Benjamini-Hochberg method. Univariable Cox’s proportional hazards regression models were used for associations with PFI, after log2 transformation and scaling of continuous measures. HR, 95%CI and p-values according to the Wald statistics were reported. For multivariable Cox’s regression, variables with a p-value < 0.1 were entered in a stepwise forward-backward model minimizing the Akaike Information Criterion (package MASS(Venables & Ripley, 2002)). “Optimal” (quoted because considerable as such only in the examined case set) cutoffs to dichotomize continuous variables were obtained using binary class labels (i.e. NR vs PR patients) and maximizing the accuracy to correctly classify those classes with the package cutpointr(Thiele, 2019). Forest plots representing adjusted HR and 95%CIs were generated with the package survminer(Kassambara *et al*, 2019). To estimate the relative abundances of cell types in gene expression mixtures from the Cancer Genome Atlas high-grade ovarian cancer data (OVCAR) - processed and normalized as described in Roelands J, et al.,(Roelands *et al*, 2019) the count matrix was analyzed using CIBERSORTx(Newman *et al*, 2019) on the dedicated web tool available at the URL https://cibersortx.stanford.edu/. The following parameters were set for the analysis: impute cell fractions, signature file LM22.update-gene-symbols.txt, batch correction enabled, batch correction mode B-mode, disable quantile normalization true, run mode absolute, permutation number 1,000. The job was performed on March 6, 2020. To derive cancer cellularity, we used ESTIMATE(Yoshihara *et al*., 2013) with default parameters, after log2 transformation and offsetting count data by a value = 1. Single sample gene set enrichment analysis for selected immune phenotypes, gene, and hallmark immune signatures was obtained as previously described(Bindea *et al*, 2013; Liberzon *et al*., 2015; Roelands *et al*., 2019; Thorsson *et al*., 2018). For univariable analysis of correlations between SLFN11 expression and cell fractions, we calculated the Spearman’s correlation coefficient. P-values were adjusted for multiple testing using the Benjamini-Hochberg method. For multivariate analysis of CIBERSORTx cell fractions and SLFN11 expression in the HGSOC TCGA dataset, we included variables with FDR < 0.05 for correlation with SLFN11, including cancer cellularity, after normalizing vectors as follows: we first removed near-zero variance variables (*caret* package(Kuhn *et al*, 2020)), then we pseudo-normalized data using the Tukey’s ladder of power transformation method (*rcompanion* package(Mangiafico, 2020)), finally we centered and scaled them. The relationship between variables was represented using a variable correlation plot, with SLFN11 expression as a supplementary quantitative variable(Lê *et al*, 2008). For survival analyses, we selected TCGA OVCAR (HGSOC) cases with the following characteristics: stage IIIc/IV, histologic grade III, and with PFI > 28 days. SLFN11 was considered “high” when in the top two tertiles of expression, and “low” otherwise. Univariable Cox’s regression and Kaplan-Meier curves were used as described above. For multivariable Cox’s regression, we first transformed bimodal CIBERSORTx cell fractions into binary factors (“present” vs. “absent”) if the Hartigan’s dip test for unimodality was rejected with p-value < 0.01. We then performed feature selection starting from stage, age, SLFN11 transcript, and CIBERSORTx variables. To do so, we fitted the Cox’s regression model by regularizing it with a lasso penalty, using the package glmnet(Friedman *et al*, 2010) with default options (α = 1, 10-fold cross-validation), and iterating it 1,000 times to obtain the minimum average error of the regularization parameter lambda for variable selection. Finally, selected variables were entered in a Cox’s multiple regression model to report HR point estimates and 95% CI. The dendrogram of similarity between immunologic signatures was built through hierarchical clustering using the Ward’s criterion agglomeration method and Euclidean distance between variables(Murtagh & Legendre, 2014).

### Power considerations

The sample size for the present study was meant to identify a clinically significant difference in SLFN11 expression between platinum-resistant (PR) patients, defined as relapsing within six months from the end of chemotherapy, and platinum-sensitive (PS) ones (i.e. relapsing beyond six months from the end of treatment). The suggested size of the collected cohort was based on our previous findings of a very significant hazard ratio in terms of overall survival between “SLFN11-high” and “SLFN11-low” HGSOC patients, with SLFN11 levels deemed so if being above or below the median for the considered cohort(Zoppoli *et al*., 2012). The required sample size would be of 24 patients, equally allocated in two groups of 12 PR and 12 PS ones, assuming a proportion of “SLFN11-high” patients of 10% in the first group and 70% in the second groups, with two-tailed α = .05 and 1 - β = .9 (z test family, G*Power 3.1.4). Assuming that 20% of samples could not be analyzed due to failure in sample processing or testing, it was estimated that 28 samples were a sufficient number needed to test the aforementioned hypothesis.

## Acknowledgements

Maria Udriste, Sophie E. Willis and Elisabeth Wiseman are thanked for generating sections of FFPE samples for IHC analyses. G.Z. wishes to thank Dr. P. Blandini for his precious insights throughout the presented research.

## Author contributions

Conceptualisation & design: C.W., D.B., G.Z; Methodology, software & data collection: C.W., F.D., A.G., F.G., J.R., L.R., D.B., G.Z.; Writing - original draft and visualisation: C.W., D.F., D.B., E.L., G.Z.; Writing -review & editing: all authors; Supervision: E.L., G.Z.; Project administration: N.C., I.RC., E.L., G.Z; Funding acquisition: E.L., G.Z.

## Supplementary figure legends

**Figure S1:**
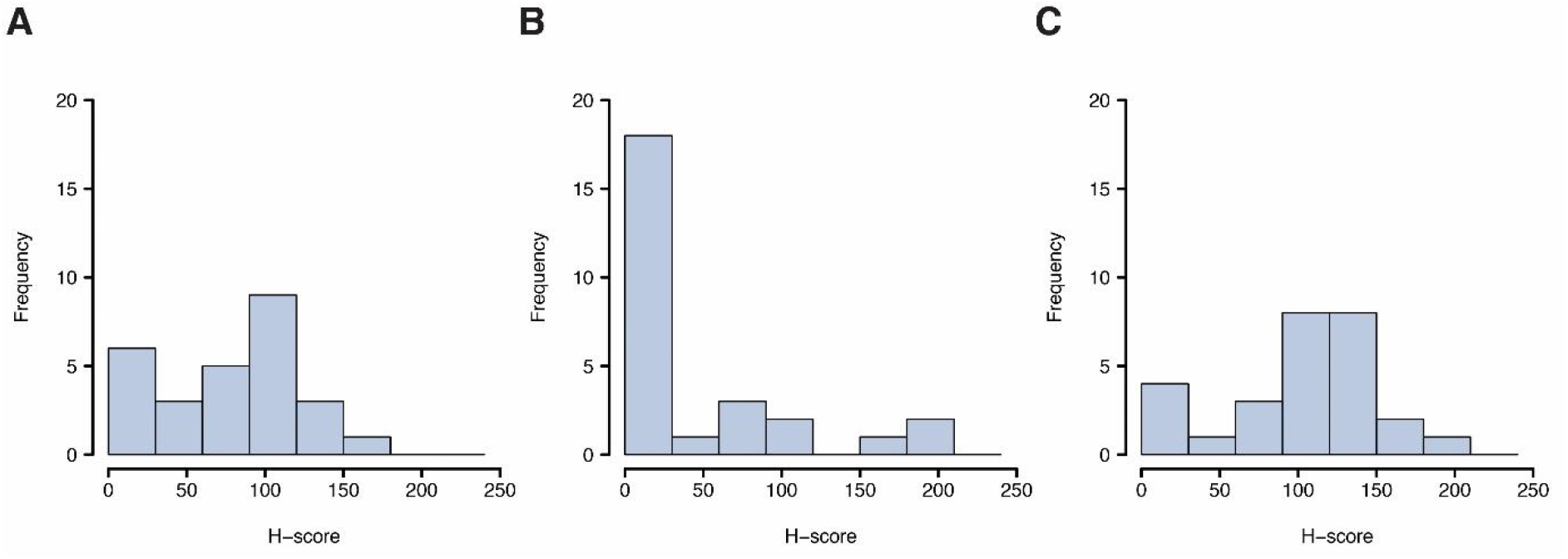
Frequency distribution of SLFN11 H-scores in the analyzed HGSOC case set. Nuclear SLFN11 protein in cancer and non-cancer cells was blindly assessed with automated image analysis by Halo and quantified as an H-score in all nuclear cells (panel A), cancer cells (panel B), and non-cancer cells (panel C). *y* axis: frequency of samples within each H-score bin; *x* axis: H-score values, subdivided into increasing 30-unit bins.

**Figure S2:**
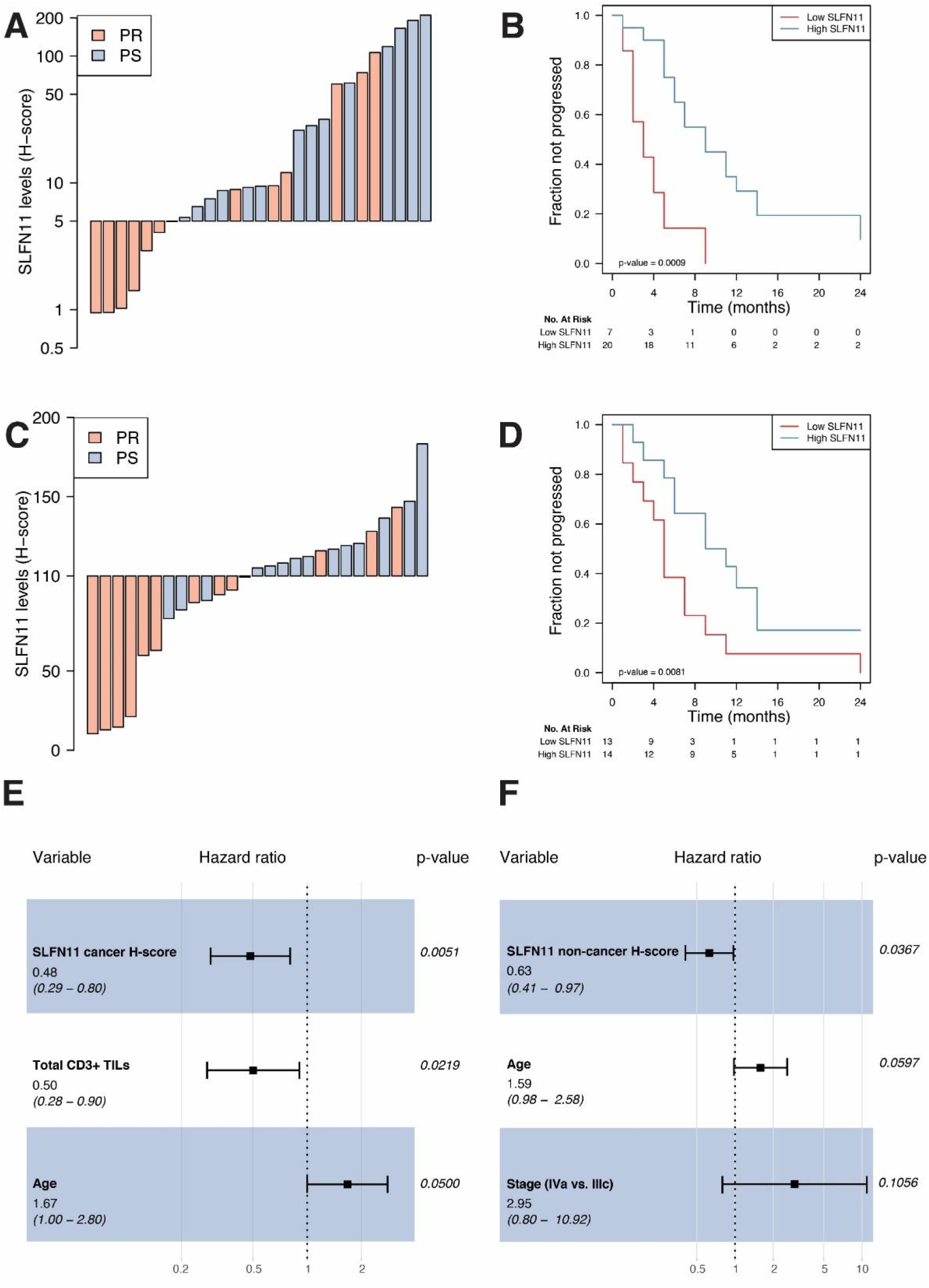
SLFN11 is independently prognostic in HGSOC also when assessed in cancer or non-cancer cells only. ***Panels A and C:*** Waterfall plot showing SLFN11 protein levels in cancer (panel A) and non-cancer (panel C) cells, colored by platinum sensitivity: SLFN11 protein is reported as H-score (*y* axis), whereas cases are reported by increasing values (*x* axis) and colored in red if platinum-refractory (PR) or light blue if non-refractory (NR). ***Panels B and D:*** Kaplan-Meier plots showing progression-free interval (PFI) stratified by SLFN11 cancer protein levels (“high” if H-score > 5, “low” if < 5, panel B) and non-cancer protein levels (“high” if H-score > 110, “low” if < 110, panel B). The progressed fraction of patients (*y* axis) is plotted against time expressed in months from the end of first-line chemotherapy, censored at 24 months (*x* axis). Numbers at risk are reported below the plots. P-values in the bottom left of the plots are from the Wald statistics for the univariable Cox’s regression. ***Panels E and F:*** Forest plots of hazard ratios (x axis, in log scale) for SLFN11 protein in cancer (panel E) and non-cancer (panel F) cells together with variables retained by the multivariable model generated for overall SLFN11. Point HR estimates are reported below each variable together with 95% confidence intervals (95%CI) in parentheses, whereas adjusted p-values for each variable are on the right side of the plot. Filled black squares represent HR estimates, with relative 95%CI shown as horizontal lines with brackets.

**Figure S3:**
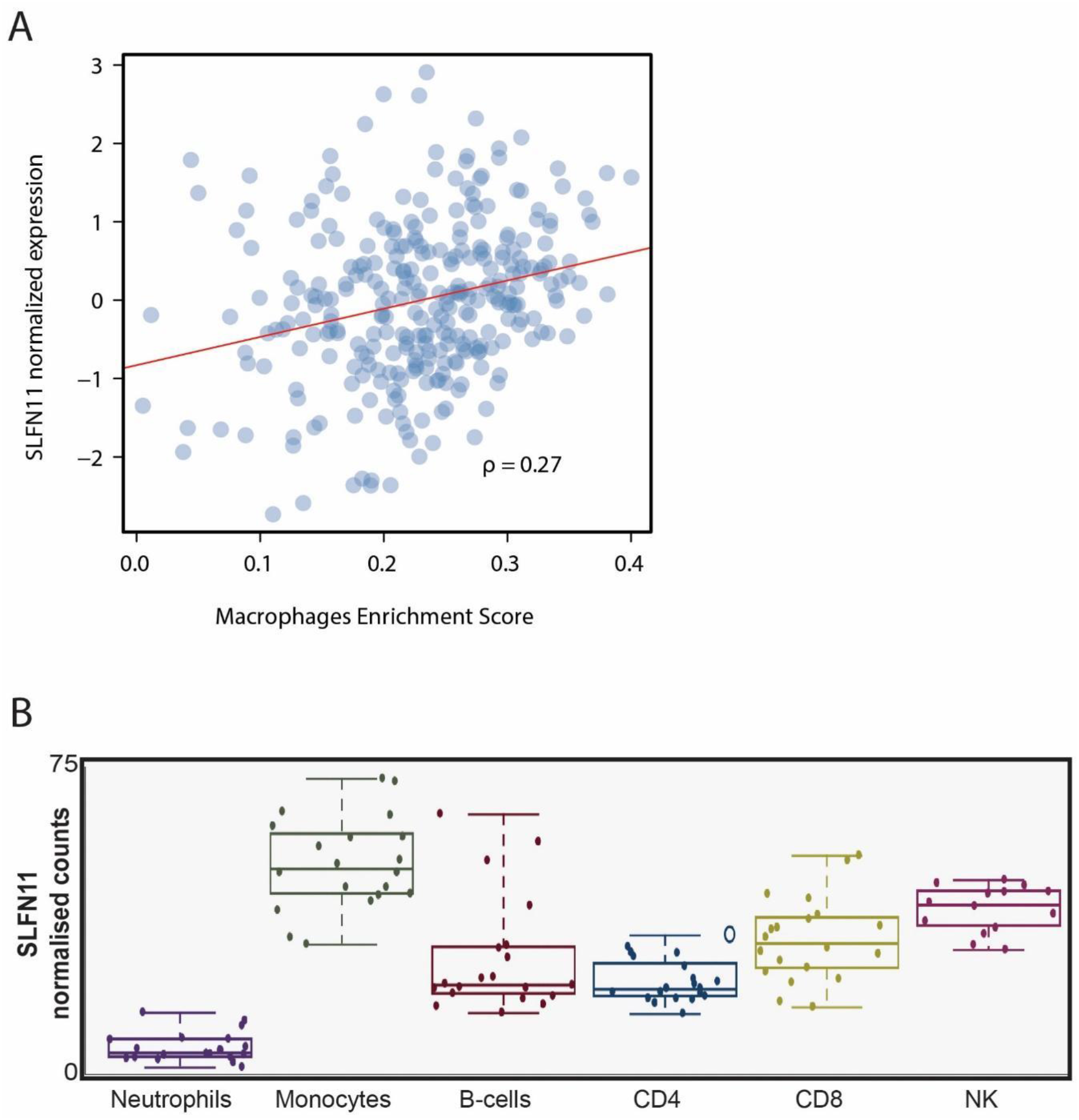
SLFN11 is expressed in a subset of leukocyte subpopulations. ***Panel A:*** Scatterplot representing SLFN11 expression (log-transformed normalized counts, *y* axis) as a function of GSEA macrophage enrichment score (ES, *x* axis) in TCGA OVCAR dataset (N = 302); ρ is the Spearman’s correlation coefficient, the least squares regression are represented by the red lines, whereas dots are measurements of SLFN11 expression by ES in individual samples. ***Panel B:*** Box plots of publicly available RNA-sequencing results (GEO accession number GSE60424) for SLFN11 in sorted leukocyte subpopulations from patients. NK: natural killer cells. GEO: gene expression omnibus.

**Figure S4:**
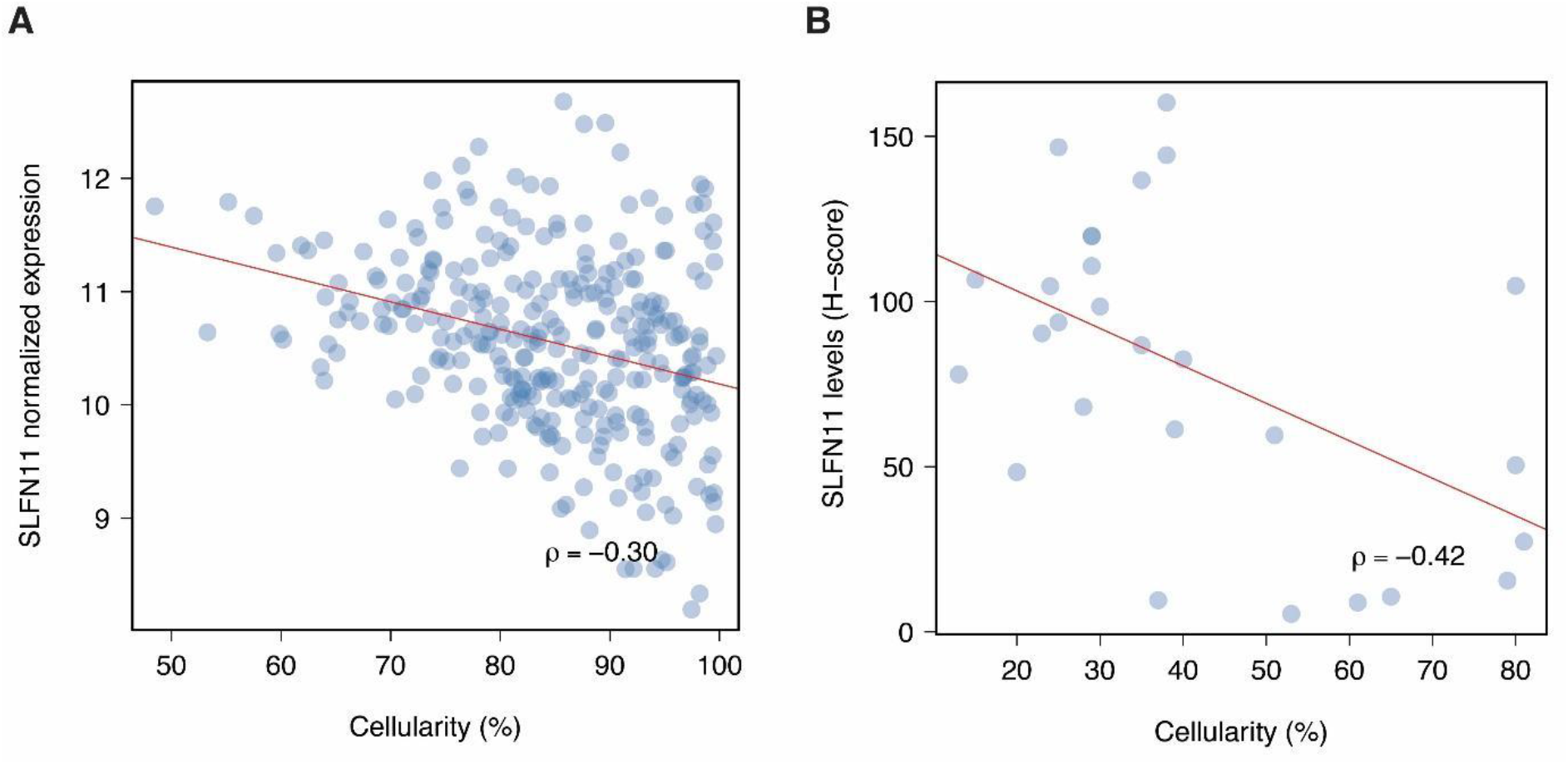
Cancer cellularity is negatively correlated with SLFN11 in ovarian cancer. ***Panel A:*** Scatterplot representing SLFN11 transcript (log-transformed normalized counts, *y* axis) as a function of cancer cellularity inferred using ESTIMATE (*x* axis) in TCGA OVCAR dataset (N = 302). ***Panel B:*** Scatterplot representing SLFN11 protein measured in all nuclear cells (overall H-score, *y* axis) as a function of cancer cellularity measured by HALO (*x* axis) in our cohort (N = 27). ρ is the Spearman’s correlation coefficient, the least squares regression are represented by the red lines, whereas dots are measurements of SLFN11 protein levels by cancer cellularity in individual samples.

**Figure S5:**
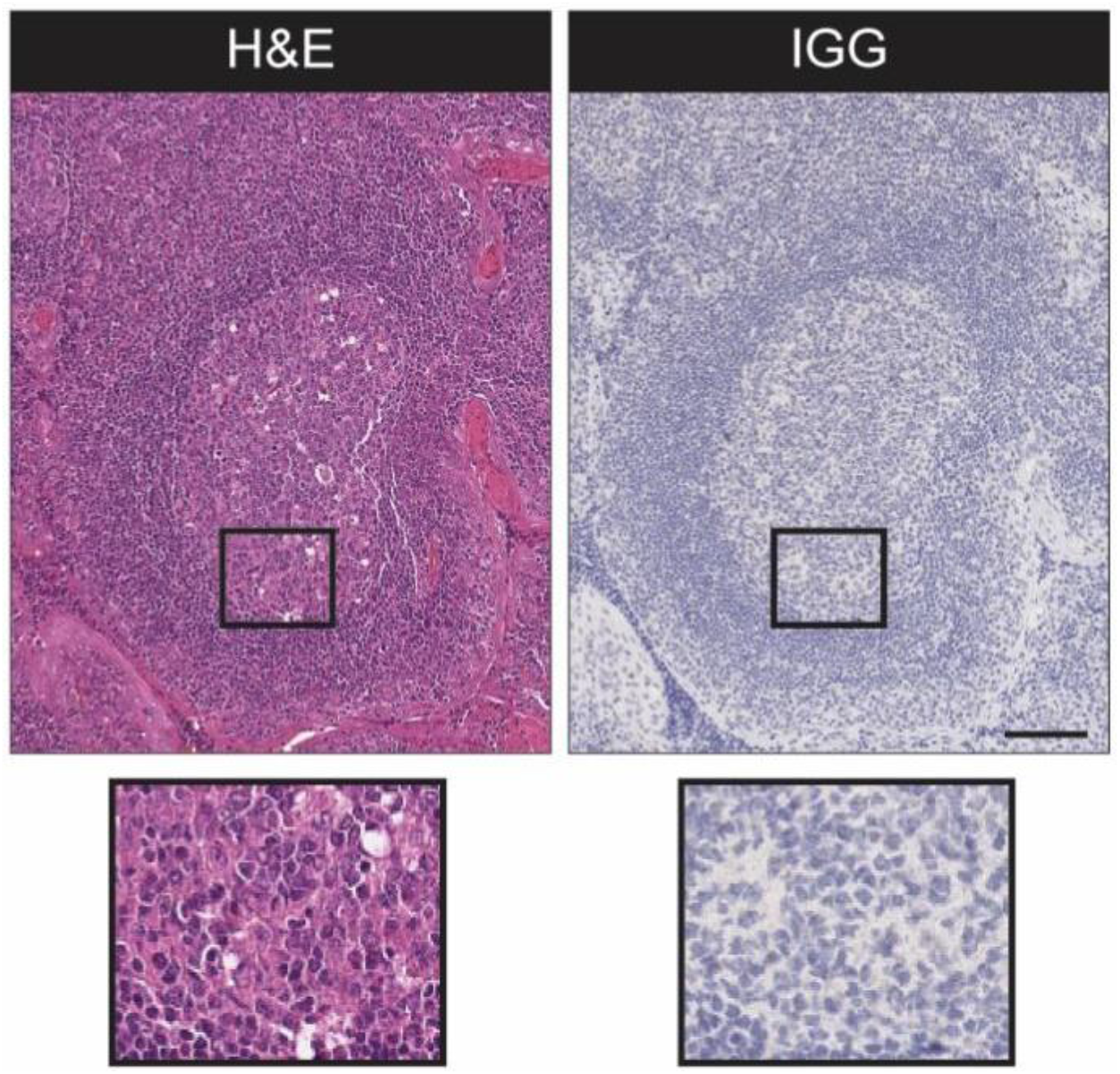
SLFN11 in tonsil tissues. Representative images of IGG IHC (negative control) and serial sections of tonsil tissue. Shown is a lymphoid follicle with the oval-shaped germinal center, the surrounding mantle zone and, at the outer layer of the lymphoid follicle, the paracortical zone. The insets show H&E stained tonsil tissue and confirm the absence of signal in the negative control sample.

